# Molecular basis of enfumafungin inhibition and resistance in *Candida glabrata* β-1,3-Glucan synthase FKS1

**DOI:** 10.64898/2026.02.07.704550

**Authors:** Ping Yang, Chu Qi, Shuchang Du, Xinlin Hu, Zhengkang Hua, Yizheng Yang, Di Zhang, Yan Ke, Mingjie Zhang, Xiaotian Liu, Tao Chen, Min Zhang, Hongjun Yu

**Affiliations:** Department of Biochemistry and Molecular Biology, School of Basic Medicine, Tongji Medical College and State Key Laboratory for Diagnosis and Treatment of Severe Zoonotic Infectious Diseases and Hubei Key Laboratory of Natural Active Polysaccharides, Huazhong University of Science and Technology, Wuhan, China; Department of Pathogen Biology, School of Basic Medicine, Tongji Medical College and State Key Laboratory for Diagnosis and Treatment of Severe Zoonotic Infectious Diseases, Huazhong University of Science and Technology, Wuhan, China; School of Life Sciences, Southern University of Science and Technology, Shenzhen, Guangdong, China; Department of Infectious Diseases, Tongji Hospital, Tongji Medical College and State Key Laboratory for Diagnosis and Treatment of Severe Zoonotic Infectious Disease, Huazhong University of Science and Technology, Wuhan, Hubei Province, China; Cell Architecture Research Center, Huazhong University of Science and Technology, Wuhan, China

## Abstract

β-1,3-Glucan synthase FKS1 is essential for fungal cell wall biogenesis and serves as a validated target of widely prescribed antifungal drugs. However, the molecular mechanism of pathogenic FKS1, its modes of inhibition, and the associated resistance mechanisms remain elusive, hindering the antifungal development. Here, we focus on *Candida glabrata* FKS1, a clinically relevant target frequently linked to antifungal resistance. We present cryo-EM structures of *C. glabrata* FKS1 in multiple detergent environments and in complex with the triterpenoid antifungal enfumafungin. Integrated functional studies revealed a key catalytic residue within the conserved ED motif and uncovered structural adaptations to different membrane-mimetic environments, rationalizing FKS1’s sensitivity to its surrounding membrane context. Enfumafungin binds at the extracellular membrane leaflet of cgFKS1 transmembrane domain, engaging a convex surface separated from the cytosolic active site, consistent with its role as a non-competitive inhibitor. Further analyses identified key enfumafungin-binding residues from TM5-TM6 and elucidated the molecular basis of drug resistance in *C. glabrata*. Collectively, these findings establish a mechanistic framework for pathogenic FKS1 function and inhibition and provide a molecular basis for the rational design of next-generation antifungal therapeutics.

## Introduction

Fungal infections constitute an escalating global health threat, affecting over one billion individuals annually^1^, with invasive mycoses responsible for approximately 3.8 million deaths each year^2^. Therapeutic options remain strikingly limited, confined to three major antifungal classes-azoles, echinocandins, and polyenes^3,4^. Within this constrained treatment landscape, *C. glabrata* has emerged as a particularly formidable threat, owing to its increasing prevalence^5^, mortality rates exceeding those of *Candida albicans*^6^, and its pronounced propensity to acquire resistance to first-line agents, especially azoles and echinocandins^7^. These characteristics have led the World Health Organization to designate *C. glabrata* as a high-priority fungal pathogen^8^.

Pathogenic fungi synthesize an essential structural component of their cell walls, β-1,3-glucan, a polysaccharide that is absent in mammalian hosts^9^. This polymer is indispensable for maintaining fungal cell integrity, growth, and virulence, and its depletion or structural disruption leads to severe cell wall defects and eventual fungal death^10–15^. The synthesis of β-1,3-glucan is catalyzed by the β-1,3-glucan synthase complex, whose core catalytic activity is mediated by the membrane-embedded FKS subunit^16–18^. This enzyme utilizes UDP-glucose as the donor substrate to elongate the glucan polymer, which is concurrently extruded across the plasma membrane (**Fig. 1a**). Owing to its fungal-specificity, essential physiological role, and high degree of conservation across pathogenic fungi, FKS represents one of the most clinically validated and therapeutically attractive targets for broad-spectrum antifungal drug development^19,20^.

**Figure 1.**
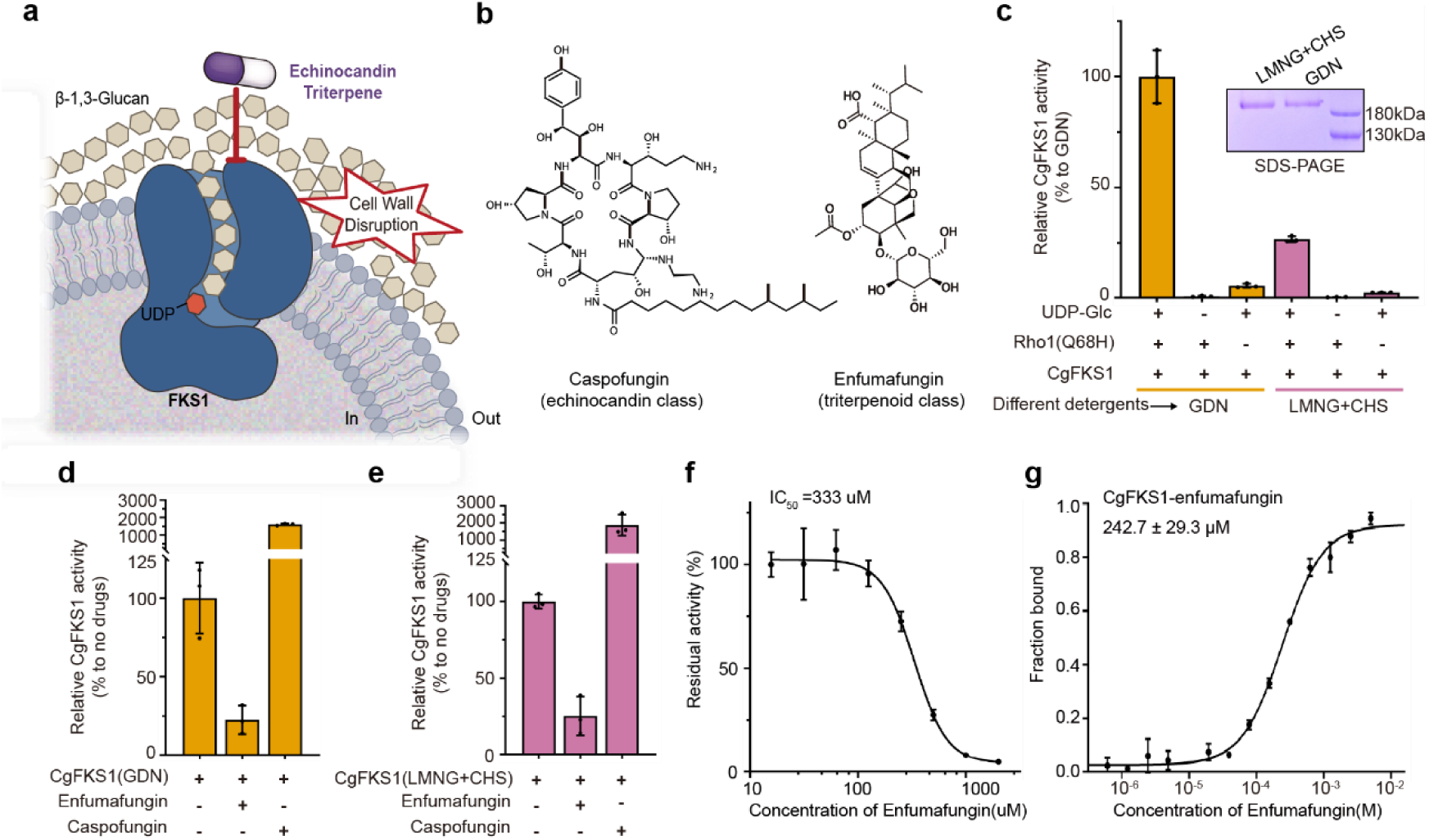
Functional characterization of *Candida glabrata* FKS1. **a**, Schematic illustration of the antifungal drug target FKS1. FKS1 catalyzes the synthesis of β-1,3-glucan from UDP-glucose and is targeted by major classes of antifungal agents, including echinocandins (for example, caspofungin) and triterpenoids (for example, enfumafungin). **b**, Chemical structures of caspofungin and enfumafungin. **c**, In vitro activity of CgFKS1 purified in either GDN or LMNG supplemented with cholesterol hemisuccinate (LMNG/CHS), measured in the presence of the indicated effectors, including UDP-glucose (UDP-Glc) and the constitutively active Rho1 variant Rho1(Q68H). Inset, SDS-PAGE analysis of cgFKS1 purified in two detergent systems. **d-e**, Effects of caspofungin and enfumafungin on the activity of cgFKS1 purified in GDN (**d**) or LMNG/CHS (**e**). **f**, Curve of cgFKS1 inhibition by enfumafungin, yielding an IC₅₀ value of ∼333 μM. Data are shown as mean ± s.d. from three independent experiments. **g**, Microscale thermophoresis (MST) analysis of the binding affinity between cgFKS1 and enfumafungin. Data represent the mean ± s.d. of three independent measurements.

Therapeutic targeting of FKS has yielded two principal classes of antifungal agents: the clinically established echinocandins and the emerging triterpenoids (**Fig. 1a-b**)^4,20–24^. Echinocandins (e.g., caspofungin, micafungin, and anidulafungin, rezafungin) represent the first successful class of FKS inhibitors approved for clinical use^25^. However, their widespread application has been accompanied by a concerning rise in resistance, particularly among *Candida* species such as *C. glabrata*^7,26^. This resistance is predominantly driven by acquired mutations within specific "hotspot" regions of the FKS genes and is strongly associated with clinical treatment failure, although the underlying resistance mechanism remains elusive^26,27^. These limitations have spurred the development of the triterpenoid antifungals. Among this class, enfumafungin exhibits potent, broad-spectrum, and durable antifungal activity against a range of pathogens, including *Aspergillus fumigatus* and *Candida species*^28–30^. Notably, ibrexafungerp, a semi-synthetic derivative of enfumafungin, was recently approved as the first oral agent in this class and demonstrates promising activity against some echinocandin-resistant strains^23,29,31,32^. Intriguingly, the genetic screening with ibrexafungerp revealed a partially overlapping resistance profile with echinocandins, underscoring the complexity of action and resistance mechanism across different classes of FKS-targeting antifungals^33–35^. Despite these advances, critical gaps remain. While recent structural studies have revealed the architecture of yeast *Saccharomyces cerevisiae* FKS1^36–38^, structural knowledge of FKS from any pathogenic fungi is entirely lacking. More importantly, despite their therapeutic potential, the precise structural and molecular mechanism of action for FKS-targeting antifungals remains poorly understood. This knowledge gap impedes a definitive understanding of inhibition and resistance mechanisms, posing a major barrier to the rational design of next-generation FKS-targeting therapeutics.

Here, we report the cryo–electron microscopy structure of the *C. glabrata* FKS1 in both apo- and enfumafungin-bound states. Combined with structure-guided functional analyses, our work defines key molecular determinants of FKS1 activity, elucidates the mechanism of enfumafungin-mediated inhibition, and provides a structural framework for understanding clinically observed drug-resistant mutations in this essential antifungal target.

## Results

### Functional characterization of *Candida glabrata* FKS1

We expressed and immunopurified *C. glabrata* FKS1 (cgFKS1) carrying a C-terminal 3×FLAG tag (see Methods). Given previous indications that the membrane environment influences FKS1 function^36,39,40^, cgFKS1 was independently purified in two commonly used detergent systems, glyco-diosgenin (GDN) and lauryl maltose neopentyl glycol supplemented with cholesterol hemisuccinate (LMNG/CHS). Size-exclusion chromatography (SEC) analysis showed that cgFKS1 exhibited well-behaved elution profiles under both purification conditions, indicative of a homogeneous preparation.

To characterize the function of purified cgFKS1, we employed our established in vitro assay that quantifies UDP production as a readout of FKS1 activity^36^. Enzymatic activity was readily detected for cgFKS1 purified in either detergent (**Fig. 1c**). Consistent with previous reports, cgFKS1 activity was strictly dependent on the small GTPase Rho1, as addition of the constitutively active Rho1(Q68H) variant strongly stimulated glucan synthase activity^41^. Notably, cgFKS1 purified in LMNG/CHS exhibited markedly reduced activity, reaching only ∼25% of that observed for the GDN-purified enzyme (**Fig. 1c**), underscoring the pronounced sensitivity of FKS1 function to its membrane-mimetic environment.

Echinocandin and triterpenoid classes are two major types of antifungal agent that targets FKS1 (**Fig. 1a-b**)^4,20–24^. We next examined the effects of antifungal compounds targeting FKS1, focusing on caspofungin and enfumafungin, the two representative members of the echinocandin and triterpenoid classes, respectively^24,30^. Strikingly, cgFKS1 purified in either GDN or LMNG/CHS exhibited comparable drug responses (**Fig. 1d-e**): enfumafungin potently inhibited enzymatic activity to below 25% of basal levels, whereas caspofungin induced a pronounced hyperactivation exceeding a tenfold increase. Dose-response analysis revealed a typical inhibitory profile, with the IC_50_ determined as 333 µM (**Fig. 1f**). To further characterize drug-cgFKS1 interactions, we performed microscale thermophoresis (MST) measurements. These analyses showed that enfumafungin clearly binds to cgFKS1 with an affinity of ∼240 μM (**Fig. 1g**). Intriguingly, we noted that caspofungin induced possible aggregation, which precluded accurate affinity determination; this may underlie the strikingly different effects of caspofungin compared with enfumafungin (**Fig. 1d-e**), awaiting futher investigation.

### Architecture of *Candida glabrata* FKS1 and its catalytic mechanism

We performed single-particle cryo-EM analysis of cgFKS1 purified in two detergent systems (GDN or LMNG/CHS) and determined their structures at overall resolution of 3.63 Å and 2.71 Å, respectively (**Supplementary Table 1**). The architecture of cgFKS1, exemplified by the structure obtained under the GDN condition, spans approximately 100 Å in width and 115 Å in height, and comprises a bilobed cytosolic catalytic head mounted on an extensive transmembrane scaffold (**Fig. 2a-b**). This organization closely resembles that of previously reported yeast FKS1 structures^36–38^, underscoring a conserved architectural framework. The cytosolic region consists of a glycosyltransferase (GT) domain (residues 700-1253) and an adjacent accessory (AC) domain (residues 132-429), which together form a compact catalytic assembly positioned above the membrane (**Fig. 2b**). The transmembrane region is composed of 17 helices arranged into two major bundles (TM1–6 and TM7–17) (**Fig. 2b-c**). Notably, the predicted TM10 is not resolved in the cgFKS1 structure and appears disordered, in contrast to its defined counterpart in yeast FKS1^36^. On the extracellular side, the transmembrane region is tightly capped by multiple interweaving luminal loops that are stabilized by an N-linked glycan at N1836 and two disulfide bonds (C644-C655 and C1315-C1332) (**Fig. 2b-c**). On the cytosolic side, the TM1-TM6 bundle is capped by the cytosolic catalytic head, whereas several loops connecting the TM7-17 bundle remain disordered, leaving the cytosolic ends of these helices exposed (**Fig. 2b-c**). Notably, helices TM5-9 and TM11-12 form an inter-bundle interface that encloses a central cavity. This cavity extends from the cytosolic side toward, but not reaching, the extracellular surface, suggesting a conserved conduit for membrane translocation of the nascent glucan polymer (**Fig. 2b**)^36^.

**Figure 2.**
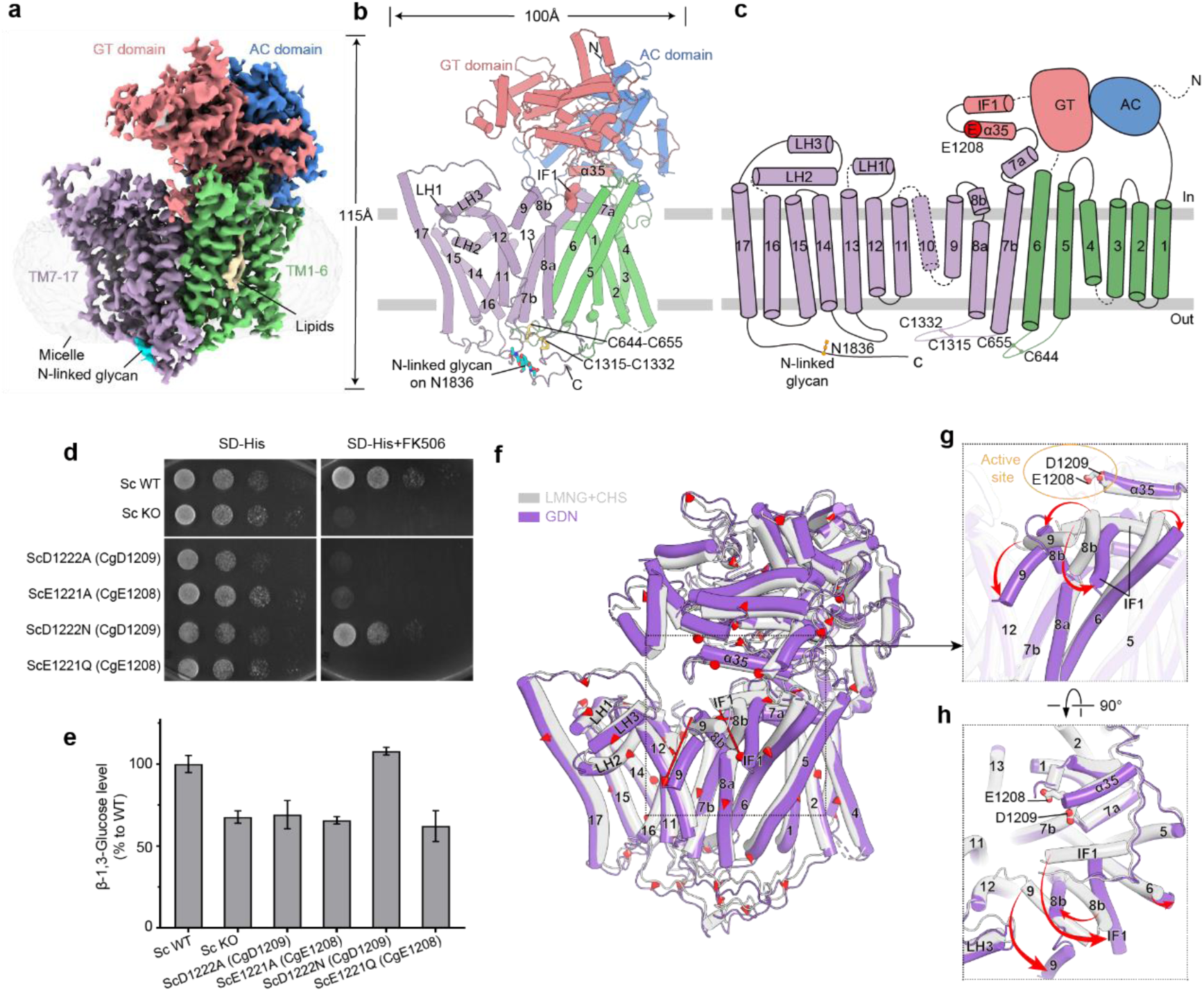
Structure of *Candida glabrata* FKS1 and its detergent-dependent conformational adaptations. **a**, Cryo-EM map of CgFKS1 in GDN, colored by domain: GT domain (red), AC domain (blue), transmembrane helices TM1–6 (green) and TM7–17 (purple). **b**, Cartoon representation of CgFKS1 structure in GDN, viewed parallel to the membrane, colored as in (**a**). Key conserved features at the active site are highlighted: the amphipathic interface helix IF1 and the adjacent helix α35 containing the catalytic ‘ED’ motif. Two conserved disulfide bonds and the N-linked glycan on conserved N1836 are indicated. **c**, Topology diagram of CgFKS1, colored as in (**a**). The catalytic residue E1208 within the ‘ED’ motif on helix α35 is marked by a red circle. Unresolved flexible regions are depicted as dashed lines. **d**, In vivo functional assay of FKS1 ED motif mutations. Growth of the indicated strains was assayed for 72 h in the absence (left) or presence (right) the FKS2-specific inhibitor FK506. Experiments were repeated three times with consistent results. **e**, Cell wall glucan levels in strains carrying the indicated FKS1 ED motfi mutations. **f**, Structural overlay of CgFKS1 structures purified in LMNG/CHS (grey) and GDN (purple), with red arrows indicating detergent-sensitive conformational changes. Major rearrangements occur near the active site, involving TM6, TM8b, TM9, and IF1. **g-h**, Two orthogonal close-up views highlighting the conformational shifts identified in (**f**) (red arrows).

Positioned immediately above this putative glucan translocation pathway, the active site of cgFKS1 resides at the interface between the TM1-6 bundle and the cytosolic GT domain (**Fig. 2b**). This site is conserved with that of yeast FKS1 and cellulose synthase BcsA and is composed of Y835, E837, K1069 and N1072 ^42^, which have been implicated in donor binding based on previous yeast studies^36^. As for the precise catalytic mechanism, inverting glycosyltransferases typically employ an aspartate or glutamate residue as the general catalytic base^43^. In FKS1, two acidic residues within the conserved ED motif have been proposed as catalytic candidates, although their precise roles have remained unsolved^36^. To interrogate the catalytic contribution of the ED motif, we exploited the genetically tractable yeast system and introduced point mutations into the chromosomal scFKS1 locus (see Methods). Specifically, we substituted scFKS1 E1221 (corresponding to cgFKS1 E1208) with alanine or glutamine, and scFKS1 D1222 (corresponding to cgFKS1 D1209) with alanine or asparagine, and assessed residual FKS1 activity using plate-based growth assays^44^ (**Fig. 2d**). Whereas the D1222A mutation impaired FKS1 function, the conservative D1222N substitution had no detectable effect (**Fig. 2d**). In contrast, both E1221A and E1221Q mutations rendered FKS1 inactive, phenocopying an FKS1 knockout strain (**Fig. 2d**). Consistently, quantification of cell wall glucan levels revealed that all mutants except D1222N exhibited a marked reduction in glucan content to levels comparable to those of the FKS1-deficient strain, whereas D1222N remained indistinguishable from wild type (**Fig. 2e**).

Together, these results support the glutamate residue in FKS1 ED motif (scFKS1 E1221; cgFKS1 E1208) as the general catalytic base required for glucan polymerization. Notably, this mechanism contrasts with that of the structurally related the bacterial cellulose synthase BcsA, where the aspartate residue of the ED motif is thought to serve as the catalytic base^42^. This functional divergence highlights mechanistic variation among processive glycosyltransferases within similar modular architectures and points to unanticipated plasticity in catalytic strategies, warranting further investigation.

### Structural adaptions to different membrane-mimetic environments

To evaluate the impact of membrane-mimetic environments on cgFKS1 structure, we compared cryo-EM reconstructions obtained in GDN and LMNG/CHS detergents. Structural superposition revealed a largely preserved global fold, accompanied by localized rearrangements, yielding an overall RMSD of 1.93 Å (**Fig. 2f**). Most pronounced conformational differences cluster around the catalytic region and involve TM9, TM8b, TM6, and the amphipathic interface helix IF1 located at the membrane-cytosol boundary (**Fig. 2g-h**). Detailed structural inspection revealed a coordinated series of conformational adaptations. Relative to the LMNG/CHS condition, the GDN-purified structure exhibits a pronounced separation between the cytosolic ends of TM8b and TM6, with TM8b undergoing a particularly large displacement (**Fig. 2g**). This separation creates a space that accommodates a near 90° rotation of IF1 (**Fig. 2h**). Concomitantly, the outward swing of TM8b propagates to TM9, driving its rotation in the same direction as IF1 and resulting in deeper membrane embedding of TM9 (**Fig. 2g**).

Notably, the repositioned IF1 lies in close proximity to helix α35, which harbors the catalytic residue E1208 (**Fig. 2g**). Thus, the extensive reorganization of IF1 and its coupled transmembrane elements appears poised to influence active-site geometry and catalytic competence. Consistent with this structural coupling, cgFKS1 purified in LMNG/CHS displays substantially reduced enzymatic activity, reaching only ∼25% of that observed for the GDN-purified enzyme (**Fig. 1c**). Together, these findings suggest a membrane-environment-dependent allosteric transition linking the transmembrane scaffold to the catalytic machinery, underscoring the regulatory role of the membrane milieu in glucan synthase function^36,39,40^.

### Enfumafungin binding to *Candida glabrata* FKS1

Echinocandins and triterpenoids constitute two major classes of clinically approved antifungal drugs that target the essential β-1,3-glucan synthase FKS1^4,20–24^. Using caspofungin and enfumafungin as representative compounds from each class, our biochemical assays show that detergent-purified cgFKS1 exhibited markedly greater sensitivity to enfumafungin (**Fig. 1d-e**), indicating its potent and selective inhibitory activity. Binding measurements further confirmed the specific association of enfumafungin with cgFKS1 (**Fig. 1g**). Motivated by these functional observations and the absence of mechanistic insights into any FKS1-targeting antifungal drugs, we sought to determine a high-resolution cryo-EM structure of cgFKS1 in complex with enfumafungin. Through systematic optimization, we found that cgFKS1 prepared under LMNG/CHS condition substantially improved drug occupancy, ultimately enabling us to resolve the complex structure at an overall resolution of 2.64 Å. The resulting reconstruction revealed well-defined density corresponding to enfumafungin (**Fig. 3a-b**). The compound binds to the extracellular side of FKS1 transmembrane domain and its glucopyranose headgroup protrudes from the membrane plane toward the extracellular space (**Fig. 3a and 3c**), indicating that membrane translocation is not required for drug action^45^. Notably, enfumafungin does not occupy a classical binding pocket; instead, it engages a convex surface of the transmembrane domain, framed by TM5-TM6 on one side and the surrounding membrane-mimetic detergent micelle on the other (**Fig. 3d-e**). In this configuration, the triterpenoid core packs against the hydrophobic surface of transmembrane helices, whereas the glucopyranose moiety extends into a more polar environment (**Fig. 3c**). Specifically, the glucopyranose group lies in proximity to R631 on TM5, while the triterpenoid scaffold is stabilized by multiple van der Waals interactions with surrounding hydrophobic residues, including A617, A620, Y624 and F625 on TM5, as well as W681 on TM6 (**Fig. 3d-e**). Among these residues, F625 adopts alternative side chain conformations, one of which directly contributes to drug engagement (**Fig. 3d-e**).

**Figure 3.**
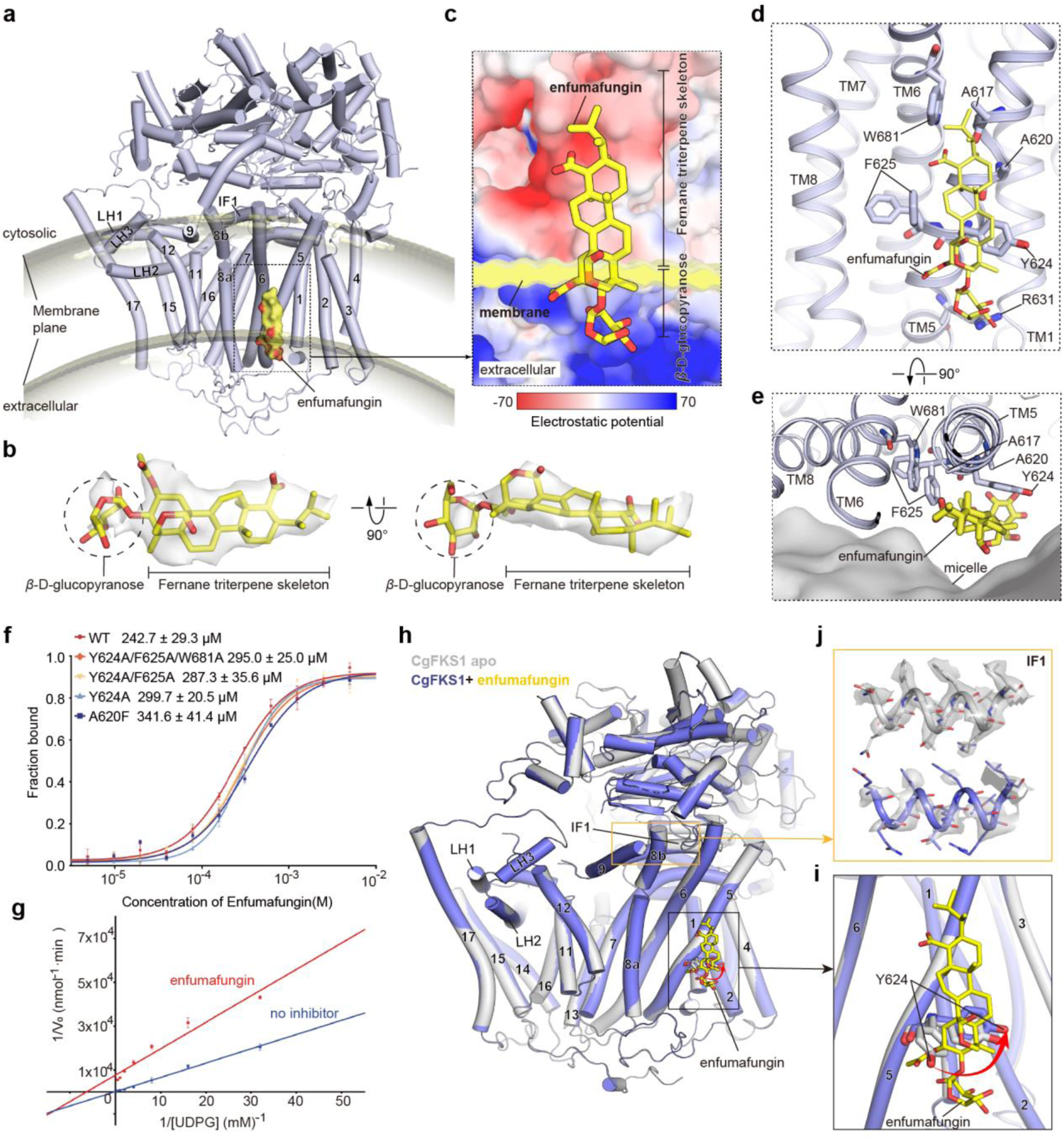
Structural basis of enfumafungin binding and inhibition of *Candida glabrata* FKS1. **a**, Cartoon representation of cgFKS1 (light blue) in complex with enfumafungin (yellow surface). The drug binds at the extracellular membrane leaflet of cgFKS1 transmembrane domain. The curved membrane plane (transparent gray) was calculated by PPM3 using a fungal plasma membrane model. Transmembrane helices TM1-17 and the interface helix IF1 are indicated. **b**, Two orthogonal views of the modeled enfumafungin superimposed with its corresponding density (transparent gray surface). **c**, Close-up view of the enfumafungin binding site, as marked by the dashed box in (**a**). cgFKS1 is rendered in electrostatic surface representation, and enfumafungin is depicted as yellow sticks. **d-e**, Close-up views of the enfumafungin interaction mode, showing key coordinating residues from TM5 and TM6. Enfumafungin and cgFKS1 residues are shown as sticks. Panel (**e**) shows a view rotated by 90° relative to (**d**), illustrating the position of the drug relative to the micelle/membrane. **f**, Microscale thermophoresis (MST) analysis of the binding affinities between cgFKS1 variants and enfumafungin. Corresponding Kd values are provided. Data are shown as mean ± SD from three independent measurements. **g**, Lineweaver-Burke plots showing the kinetic analysis of CgFKS1 (blue line) and the addition of enfumafungin (red line) by varying the amount of UDP-glucose. **h**, Structural overlay of apo (grey) and enfumafungin-bound (blue) cgFKS1. Enfumafungin binding induces local conformational rearrangements, marked by the black and yellow boxes. **i-j**, Close-up views of conformational changes associated with enfumafungin binding. **i**, In the apo state, Y624 adopts two alternative side-chain conformations, whereas enfumafungin binding stabilizes a single conformation that alleviates steric clashes with drug interactions (red arrow). **j**, Altered density of membrane-cytosol interface helix IF1, which is well ordered in the apo-state but becomes poorly resolved in the drug-bound complex.

To assess the functional relevance of this binding interface, we performed microscale thermophoresis (MST) measurements using wild-type and mutant cgFKS1 variants. Alanine substitution at Y624, F625 and W681 had minimal effects on binding affinity, whereas introduction of a bulky phenylalanine at position A620 resulted in a modest reduction in enfumafungin binding (**Fig. 3f**). These data indicate that this region contributes to ligand recognition without being strictly essential, consistent with a binding mode dominated by distributed hydrophobic interactions rather than discrete, highly specific contacts (**Fig. 3d-e**).

Kinetic analyses further detailed the inhibitory effect of enfumafungin on cgFKS1 activity. Lineweaver–Burk plots revealed that, in the presence of enfumafungin, the reciprocal initial velocity (1/ V₀) increased across the entire range of UDP-glucose concentrations tested, consistent with an overall reduction in overall catalytic efficiency (**Fig. 3g**). Concomitant changes in both the apparent Michaelis constant (K_m_) and maximal velocity (V_max_) support a noncompetitive or mixed mode of inhibition. Consistent with this kinetic behavior, the enfumafungin-binding site is located distal to the catalytic center, supporting an allosteric mechanism (**Fig. 3a**).

Structural comparison of the apo and enfumafungin-bound cgFKS1 states revealed that the overall architecture of the enzyme remains largely unchanged upon drug binding (with an RMSD of 0.16 Å), with conformational differences confined to the vicinity of the drug-binding site (**Fig. 3h**). In particular, the aromatic residue Y624 adopts two alternative side-chain conformations in the apo state, whereas enfumafungin binding stabilizes a single conformation that alleviates steric clashes and optimizes drug interactions (**Fig. 3i**). In addition, the IF1 region exhibits altered density between the two states: IF1 is well ordered in the apo structure but becomes poorly resolved in the drug-bound complex (**Fig. 3j**). IF1 contacts TM6 and the adjacent TM8 on one side and helix α35 (harboring the catalytic residue E1208) on the other (**Fig. 2g**). As we described above, the conformational plasticity of IF1, potentially influenced by the membrane-mimetic detergent environment, may underlie the observed changes in enzymatic activity (**Fig. 1c**, **Fig. 2g-h**). Moreover, the counterpart of IF1 in the bacterial cellulose synthase BcsA has been implicated in regulating the gating of cellulose translocation channel^42^. Therefore, it is conceivable that IF1 may serve as a structural bridge linking the transmembrane domain to the catalytic core, providing a plausible mechanistic basis for the noncompetitive inhibition of FKS1 by enfumafungin.

### Drug resistance mechanism

Previous genetic screens using echinocandin-class antifungal drugs have identified resistance-conferring mutations clustered within three conserved regions of FKS1^26,27,44^, termed hotspot 1 (residues 621–633 in TM5; numbering according to cgFKS1), hotspot 2 (residues 1340–1347 in TM8) and hotspot 3 (residues 677–686 in TM6) (**Fig. 4a**). Subsequent screens with ibrexafungerp, a triterpenoid derivative of enfumafungin, revealed a partially overlapping resistance landscape^7,34^, including W681 within hotspot 3; E621, F625, and I627 within hotspot 1; and A1356 in TM8, adjacent to hotspot 3 (**Fig. 4b-c**). Notably, F625 and W681 map directly to residues that engage enfumafungin in our structure (**Fig. 3d**), indicating that this binding site constitutes a focal locus for the evolution of triterpenoid resistance in fungi.

**Figure 4.**
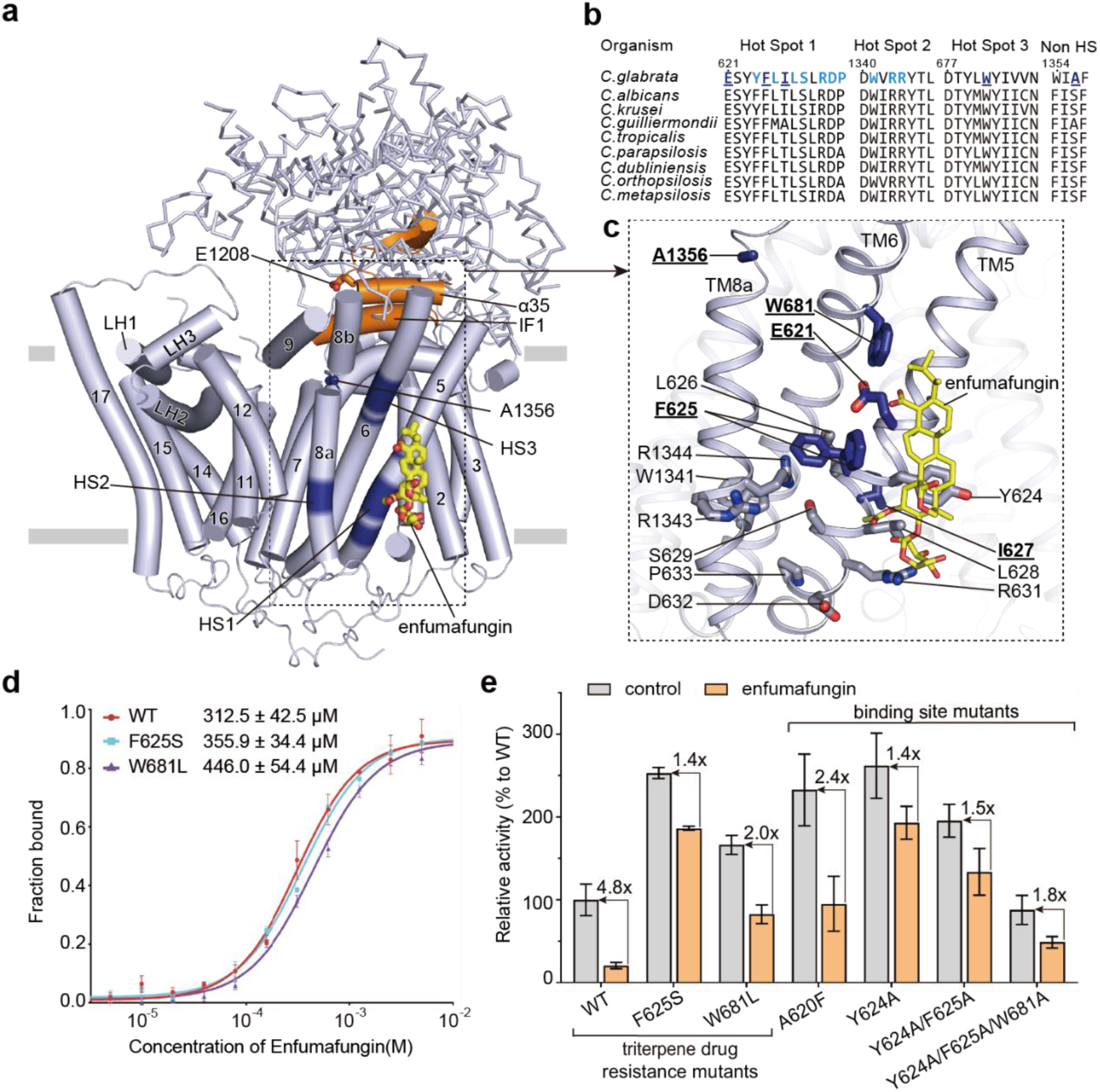
Mechanism of enfumafungin resistance. **a,** Reported echinocandin- or triterpenoid-resistance mutations from major fungal pathogens mapped onto the cgFKS1/enfumafungin complex. These mutations predominantly cluster within three resistance hotspots (blue; HS1-3), located on three transmembrane helices (TM5, TM8a, and TM6) and adjacent to the bound enfumafungin. Enfumafungin is shown as yellow sticks. Two key helices in the active site, helix α35 harboring catalytic E1208 and interface helix IF1 (see also **Fig. 2g-h**), are highlighted in orange. **b**, Sequence alignment of FKS1 across representative *Candida* pathogens, showing conservation in resistance-associated hotspot 1, hotspot 2, hotspot 3, and a non-hotspot region. Only resistance sites reported in *C. glabrata* are highlighted: triterpenoid-resistance mutations (underlined, dark blue) and echinocandin-resistance mutations (light blue). **c**, Detailed view of the drug-resistance mutation regions boxed in (**a**). Corresponding to (**b**), only the resistance-related sites characterized in *C. glabrata* are displayed as sticks: triterpenoid-resistance sites (underlined, dark blue) and echinocandin-resistance sites (light blue). **d**, Microscale thermophoresis (MST) analysis of binding affinities between enfumafungin and cgFKS1 variants, including wild-type cgFKS1 and two reported triterpenoid-resistant mutations identified as drug binding sites in the complex structure: F625S and W681L. Data are mean ± SD from three independent measurements. **e**, Enfumafungin inhibition profiles of cgFKS1 variants. Activities are normalized to wild-type cgFKS1 in the absence of enchincandin. Fold changes are indicated above the bars. Data are mean ± s.d. from three independent experiments. F625S and W681L are reported triterpenoid-resistant mutations identified as drug binding sites in the complex structure; the other variants correspond to drug binding residues identified in this study.

To interrogate the molecular basis of resistance conferred by these mutations, we generated cgFKS1 variants bearing substitutions at residues directly involved in drug engagement (W681L and F625S)^7,34^. Microscale thermophoresis (MST) measurements revealed a modest reduction in binding affinity for W681L variant, whereas the F625S substitution had minimal effect (**Fig. 4d**). Consistently, biochemical activity assays showed that both mutants exhibited reduced sensitivity to enfumafungin inhibition (**Fig. 4e**). In addition, structure-guided mutations targeting the enfumafungin-binding site (Y624A/F625A/W681A, Y624A/F625A, Y624A, and A620F) (**Fig. 3d**) similarly conferred decreased drug sensitivity (**Fig. 4e**), further supporting the functional relevance of this binding site.

### Discussions

β-1,3-Glucan synthase FKS1 is an essential enzyme for fungal cell wall biogenesis and a validated target of frontline antifungal therapies^19,20^. Despite its clinical importance, the molecular architecture of pathogenic FKS1 and the mechanisms by which antifungal agents modulate its activity have remained poorly understood. In this study, we present a comprehensive structural and functional analysis of the *C. glabrata* FKS1, capturing multiple functionally distinct states: an enfumafungin-bound state and two apo-conformations stabilized by different membrane-mimetic environments. Our results provide long-sought mechanistic insights into FKS1 catalysis and regulation, as well as the molecular basis of clinically relevant antifungal action and resistance.

A major conceptual advance of this work is the determination of a high-resolution structure of pathogenic cgFKS1 in complex with the triterpenoid antifungal enfumafungin. This provides, to our knowledge, the first direct structural insight into the mechanism of FKS1-targeting drugs. Enfumafungin binds at the extracellular membrane leaflet, engaging a convex surface formed by TM5 and TM6 rather than a buried pocket (**Fig. 3a-e**). The interaction is dominated by distributed hydrophobic contacts, with the glucopyranose moiety remaining solvent-exposed on the extracellular side (**Fig. 3c-d**). This indicates that membrane penetration is not required for inhibition, consistent with prior studies defining the action mode of FKS1-targeting antifungals^45^. In line with this binding architecture, kinetic analyses reveal a non-competitive mode of inhibition (**Fig. 3g**), supporting an allosteric mechanism in which drug binding reduces catalytic efficiency without directly occluding the active site. This is further supported by the spatial separation between the extracellular drug-binding site and the cytosolic catalytic center (**Fig. 3a**). Notably, the enfumafungin-binding site resides within transmembrane elements, including TM6, which appear to contribute to membrane-dependent allosteric regulation of FKS1 (**Fig. 2f**), a spatial relationship consistent with the allosteric nature of drug action (discussed below).

This model is reinforced by both prior and present observations linking FKS1 activity to its membrane environment. Previous studies have shown that alterations in the lipid microenvironment modulate echinocandin activity^46,47^, and we consistently observed that the functional effects of caspofungin-either inhibitory or stimulatory-depend on the detergent system used^36^. cgFKS1 prepared in LMNG/CHS exhibits markedly reduced activity compared to that purified in GDN (**Fig. 1c**). Notably, cgFKS1 is activated by caspofungin under both detergent conditions, mirroring the behavior of yeast FKS1 purified in GDN, whereas yeast FKS1 prepared in CHAPS is inhibited by the same drug^36^. The reproducibility of these activity patterns across detergents and species indicates that FKS1 activity is intrinsically sensitive to its surrounding membrane context, consistent with earlier studies linking β-1,3-glucan synthesis to defined lipid environments^39,40^.

This functional plasticity is also reflected at the structural level. Although cgFKS1 adopts a similar overall architecture across different detergent systems, we identify local yet coordinated conformational rearrangements, most prominently involving TM6, TM8b, TM9, and the amphipathic interfacial helix IF1 (**Fig. 2f-h**). IF1 is positioned at the membrane-cytosol interface, where it contacts both the transmembrane scaffold and the catalytic helix α35 containing the essential glutamate (E1208). Notably, TM6 is directly involved in enfumafungi binding (**Fig. 3d-e**). Together, these elements delineate a plausible conduit for membrane-responsive allosteric coupling between the transmembrane domain and the cytosolic catalytic core. This provides a structural framework for understanding how changes in the membrane environment and drug binding converge to regulate FKS1 activity, a model that awaits further investigation.

In addition to triterpenoids such as enfumafungin, echinocandins constitute another major class of FKS1-targeting drugs. Echinocandins are lipopeptides with differently configured lipid tails, yet a substantial subset of mutations conferring echinocandin resistance overlaps with, or lies adjacent to, those selected by triterpenoid derivatives such as ibrexafungerp (**Fig. 4b-c**). The partial overlap of these resistance determinants, together with the retained activity of triterpenoids against some echinocandin-resistant strains^31,34^, suggests that these different drug classes engage closely neighboring but not entirely identical regions of FKS1. Further studies in this direction may help clarify the molecular basis of cross-resistance and uncover shared vulnerabilities that could be exploited in future antifungal drug design.

Overall, this study advances our mechanistic and structural understanding of pathogenic (*Candida glabrata*) FKS1 function, and its inhibition by antifungal drug, providing a foundation for the rational design and optimization of the antifungal drugs targeting this enzyme.

## ACKNOWLEDGMENTS

We thank the Center for Biological Cryo-EM, Huazhong University of Science and Technology and thank Chuang Liu, Zengru Li and Hua Li for their assistance in data collection. We thank all staff members of the Cryo-EM Center, Southern University of Science and Technology for their assistance in data collection. This work was supported by the National Natural Science Foundation of China (92478121 to H.Y.), the Natural Science Fund for Distinguished Young Scholars of Hubei Province (2024AFA044 to M.Z.), Independent project of State Key Laboratory for Diagnosis and Treatment of Severe Zoonotic Infectious Disease (2025ZZ10002 to M.Z.), the National Science Foundation of China (92053112, 31971148 to H.Y., 32100575 to M.Z., 0204K0010 to X.H.), Shenzhen Medical Research Fund (A2303054 to X.L., B2302039 to M.J.Z. and X.L.), the Fundamental Research Funds for the Central Universities (5003510141, 335003510056 to H.Y. and 5003510112 to M.Z.) and Postdoctoral Fellowship Program of China Postdoctoral Science Foundation (GZB20240245 to X.H.).

## AUTHOR CONTRIBUTIONS

M.Z. and H.Y. conceived the project. M.J.Z., X.L., T.C., M.Z., and H.Y. designed experiments. P.Y. and C.Q. performed most of the experiments. S.D., X.H. Z.H., Y.Y., D.Z., and Y.K. helped with the sample preparation, functional experiments and data collection. P.Y., C.Q., X.L., T.C., M.Z. and H.Y. analyzed the data. M.Z. and H.Y. wrote the manuscript with the help of all the authors.

## DATA AVAILABILITY

The cryo-EM maps and atomic coordinates of cgFKS1-enfumafungin complex, cgFKS1 apo in GDN, cgFKS1 apo in LMNG/CHS and have been deposited and are available in the EMDB (under accession codes EMDB-aaaa, EMDB-bbbb and EMDB-cccc) and in the PDB (under accession codes xxxx, yyyy, and zzzz), respectively.

## MATERIALS AND METHODS

### The expression and purification of *Candida glabrata* FKS1 (cgFKS1)

The *Candida glabrata* FKS1 (cgFKS1) gene was cloned and expressed in the *Saccharomyces cerevisiae* expressing strain. For the purification of cgFKS1 in glyco-diosgenin (GDN) condition, cells were collected by centrifugation and lysed using a French press in lysis buffer containing 25 mM HEPES (pH 7.4), 150 mM NaCl, and protease inhibitors (cOmplete EDTA-free protease inhibitor cocktail, Roche). Cell debris was removed by centrifugation at 15,000 × g for 30 min, and membrane fractions were pelleted by ultracentrifugation at 100,000 × g for 1 h. The membrane pellet was resuspended and solubilized for 2 h at 4 °C with gentle agitation in solubilization buffer consisting of 25 mM HEPES (pH 7.4), 500 mM NaCl, 10% (v/v) glycerol, 1.5% (w/v) n-dodecyl-β-D-maltopyranoside (DDM; Anatrace), 0.15% (w/v) cholesteryl hemisuccinate Tris salt (CHS; Anatrace), 2 mM MgCl₂, and protease inhibitors (Roche). Insoluble material was removed by centrifugation at 15,000 × g for 30 min, and the clarified supernatant was applied to Anti-FLAG M2 affinity resin (Sigma) at 4 °C. After binding, the resin was washed with wash buffer containing 25 mM HEPES (pH 7.4), 150 mM NaCl, 2 mM MgCl₂, and 0.04% (w/v) GDN (Anatrace). The target proteins were eluted using the same buffer supplemented with 150 µg ml⁻¹ 3×FLAG peptide. The eluate was concentrated and further purified by size-exclusion chromatography (SEC) on a Superose 6 Increase 10/300 GL column (Cytiva) equilibrated with 25 mM HEPES (pH 7.4), 150 mM NaCl, 2 mM MgCl₂, and 0.02% (w/v) GDN. Monodisperse peak fractions were pooled, concentrated, and used for cryo-EM grid preparation, biochemical and biophysical assays. For the purification of cgFKS1 in LMNG/CHS (LMNG, lauryl maltose neopentyl glycol; CHS, cholesterol hemisuccinate) condition, similar purification procedure was applied as described above. Detergent exchange was employed throughout affinity purification and size-exclusion chromatography.

### Expression and purification of Rho1 Q68H

The construct containing N-terminal 6×His-SUMO-tagged *Saccharomyces cerevisiae* Rho1 Q68H was transformed into *Escherichia coli* BL21(DE3) cells. Transformed cells were cultured in LB medium supplemented with 100 µg ml⁻¹ ampicillin at 37 °C until the OD₆₀₀ reached approximately 0.6. Protein expression was induced by adding 0.5 mM isopropyl-β-d-1-thiogalactopyranoside (IPTG), and cultures were subsequently incubated at 18 °C for 20 h. Cells were harvested by centrifugation, resuspended in lysis buffer containing 25 mM HEPES (pH 7.4), 300 mM NaCl, and 2 mM MgCl₂, and disrupted by French press. The lysate was clarified by centrifugation, and the supernatant was applied to nickel–nitrilotriacetic acid (Ni–NTA) affinity resin. After washing, the target proteins were eluted and dialyzed into buffer composed of 25 mM HEPES (pH 7.4), 150 mM NaCl, and 2 mM MgCl₂. During dialysis, 6×His-tagged Ulp1 protease was added at a 1:200 Ulp1:Rho1 (w/w) ratio to remove the 6×His–SUMO tag. The cleaved 6× His-SUMO tag and the 6×His-tagged Ulp1 protease were removed by another run of Ni–NTA affinity chromatography. The flow-through containing untagged Rho1 Q68H was collected, concentrated, and further purified by size-exclusion chromatography on a Superdex 200 Increase column (Cytiva) equilibrated with 25 mM HEPES, (pH 7.4), 150 mM NaCl, and 2 mM MgCl_2_. The central fractions of the monodisperse peak were collected and concentrated to 4 mg ml^−1^.

### Activity Assays of cgFKS1

The activity of cgFKS1 and its variants were assayed using the UDP-Glo™ Glycosyltransferase Assay kit (Promega), following our previously established protocol^36^. Briefly, 1 µl of each purified cgFKS1 variant was added to a 30 µl reaction mixture containing 50 mM Tris-HCl (pH 7.4), 33% (v/v) glycerol, 1 mM EDTA, 6 µg ml⁻¹ Rho1-Q68H, 0.2% (w/v) CHAPS, 0.04% (w/v) CHS, and 20 mM potassium fluoride. The reaction was initiated by addition of UDP-glucose (0.25 mM final concentration). The reaction was carried out at 30 °C for 1 h. Luminescence was measured using a PheraStar FS microplate reader (BMG Labtech). For the characterization of drug inhibition, the cgFKS1 variant was first incubated with the specified concentration of caspofungin or enfumafungin at room temperature for 10 minutes, after which the remaining reaction components were added.

### Cryo-EM grid preparation and data collection

For single-particle cryo-EM analysis, 3 µL of protein sample at a concentration of 5-6 mg/ml was applied to glow-discharged Quantifoil R1.2/1.3 Au 300-mesh grids. Grids were blotted for 3 s at 4 °C and 100% humidity, and subsequently plunge-frozen in liquid ethane using an FEI Vitrobot Mark IV. For the enfumafungin-incubated sample, cgFKS1 at the concentration of 5 mg/ml was incubated with 2 mM enfumfgungin for 5 h before freezing the cryo-EM grids.

Cryo-EM data were collected on a Titan Krios transmission electron microscope (Thermo Fisher) operated at 300 kV, equipped with a Gatan K3 Summit direct electron detector and a GIF Quantum energy filter (slit width of 20 eV). Automated data acquisition was performed using EPU software (Thermo Fisher). Movies were recorded under super-resolution counting mode with a calibrated physical pixel size of 0.92 Å or 0.93 Å. Data were collected with a defocus range of −1.1 to −2.8 µm, with a total accumulated electron dose of 50 e⁻ Å⁻².

### Data processing and 3D reconstruction

The cryo-EM data processing was performed using CryoSPARC (v4.7.0)^48^. Dose-fractionated movie stacks were aligned and dose-weighted using Patch Motion Correction. Contrast transfer function (CTF) parameters for individual micrographs were estimated using Patch CTF Estimation.

For the dataset of cgFKS1 prepared in LMNG/CHS, 8,581 micrographs were obtained in total. After removing micrographs with a CTF fitting resolution worse than 6 Å, an initial set of particles from ∼1500 micrographs were picked using blob picker of cryoSPARC to generate two-dimensional (2D) class averages, which were subsequently used as templates for template particle picking. This procedure resulted in a total of 3,543,984 particles. These particles were extracted with 4 × binning and subjected to multiple rounds of 2D classification to remove poorly defined or contaminating particles. An initial ab initio three-dimensional (3D) reconstruction was generated and used as initial references for heterogeneous refinement. A cleaned subset of particles was selected and re-extracted without binning. These particles were then subjected to iterative rounds of ab initio reconstruction and heterogeneous refinement. A final subset of 577,248 particles was selected for non-uniform refinement, yielding a reconstruction with an overall resolution of 2.71 Å.

For the dataset of enfumafungin-incubated cgFKS1 prepared in LMNG/CHS condition, a total of 10,242 movie stacks were processed similarly and the final reconstruction map with an overall resolution of 2.64 Å from 548,915 particles was obtained.

For the dataset of cgFKS1 prepared in GDN condition, a total of 5,376 movie stacks were processed similarly and the final reconstruction map with an overall resolution of 3.63 Å from 155,553 particles was obtained.

For all datasets, the reported overall resolutions were determined using the gold-standard Fourier shell correlation (FSC) criterion at 0.143 ^49^. Local resolution distributions were estimated using cryoSPARC.

### Model building and refinement

The atomic model of cgFKS1 in LMNG/CHS condition was first built based on its corresponding high-resolution map (2.71 Å). Specifically, an initial model was generated using AlphaFold2 and rigid-body fitted into the cryo-EM map via Chimera^50,51^. The model was then manually adjusted and rebuilt in Coot^52^, guided by the clearly resolved densities for transmembrane helices and characteristic density features around bulky residues such as Trp, Tyr, Phe and Arg. Real-space refinement against the cryo-EM map was performed in PHENIX with secondary structure and geometry restraints applied^53^. The resulting model was used as the starting model for building the structures of cgFKS1/enfumafungin in LMNG/CHS and cgFKS1 in GDN, following a similar workflow. All final models were assessed using MOLPROBITY^54^. Structural figures were prepared using Chimera, ChimeraX, and PyMOL (Schrödinger, LLC.)^55^. Statistics for 3D reconstructions and model refinements are provided in **Supplementary Table 1**.

### Spot growth assay

Yeast strains carrying chromosomal FKS1 mutations and an FKS1 knockout (KO) strain were generated by homologous recombination method^56^. Transformants were selected on SD-His medium (Yeast Synthetic Drop-out Medium without Histidine, cat no. S0020, Solarbio) and verified by genomic PCR and DNA sequencing. For the analysis of growth phenotype, exponentially growing cells cultured in YPD medium at 30 °C with shaking at 200 r.p.m. were adjusted to an OD_600_ of 0.1. Then 4µl portions of ten-fold serial dilution were spotted onto SD-His plates with or without 1 µg ml⁻¹ FK506. Plates were incubated at 30 °C for the indicated times and imaged using a TANON 5200CE imaging system. All experiments were repeated three times with similar results.

### Quantitative determination of β-1,3-glucan in cell walls

The levels of β-1,3-glucan were determined using the aniline blue assay, as previously described^36,57^. Testing strains were cultured in YPD medium at 30 °C to exponential phase (OD_600_ = 0.5). Equal numbers of cells from each strain were harvested by centrifugation at 5,000 × g and washed twice with TE buffer (10 mM Tris-HCl, pH 8.0; 1 mM EDTA). Cell pellets were resuspended in 0.5 mL TE buffer and mixed with 0.1 mL of 6 M NaOH. Samples were incubated at 80 °C for 30 min to solubilize β-1,3-glucan. Subsequently, 2.1 mL of aniline blue solution (0.03% aniline blue, 0.18 M HCl, and 0.49 M glycine-NaOH, pH 9.5) was added to each sample. After brief vortexing, samples were incubated at 50 °C for 30 min, followed by an additional 30 min incubation at 24 °C. Fluorescence was measured using a CLARIOstar Plus plate reader (BMG Labtech) with excitation at 400 nm and emission at 460 nm (455 nm cutoff). All measurements were performed in triplicate.

---

**Supplementary Table 1.** Cryo-EM data collection, refinement and validation statistics.

|  | cgFKS1 in<br>LMNG/CHS<br>(EMDB-aaaa)<br>(PDB xxxx) | cgFKS1 in<br>GDN<br>(EMDB-bbbb)<br>(PDB yyyy) | cgFKS1 with<br>enfumafungin<br>(EMDB-cccc)<br>(PDB zzzz) |
| --- | --- | --- | --- |
| **Data collection and processing** |  |  |  |
| Magnification | 130,000 | 130,000 | 130,000 |
| Voltage (kV) | 300 | 300 | 300 |
| Electron exposure (e-/Å<sup>2</sup>) | 50 | 50 | 50 |
| Defocus range (µm) | -1.1 to -2.8 | -1.1 to -2.8 | -1.1 to -2.8 |
| Pixel size (Å) | 0.93 | 0.92 | 0.93 |
| Symmetry imposed | C1 | C1 | C1 |
| Initial particle images (no.) | 3,543,984 | 2,753,707 | 4,814,902 |
| Final particle images (no.) | 577,248 | 155,553 | 548,915 |
| Map resolution (Å) | 2.71 | 3.63 | 2.64 |
| FSC threshold | 0.143 | 0.143 | 0.143 |
| **Refinement** |  |  |  |
| Initial model used (PDB code) | AlphaFold2 | This study | This study |
| Model resolution (Å) | 2.71 | 3.63 | 2.64 |
| FSC threshold | 0.5 | 0.5 | 0.5 |
| Model composition |  |  |  |
| Non-hydrogen atoms | 12,354 | 11,658 | 12,398 |
| Protein residues | 1487 | 1407 | 1487 |
| Ligands | 8 lipids; 1 N-glycan | 8 lipids; 1 N-glycan | 8 lipids; 1 N-glycan; 1 enfumafungin |
| *B* factors (Å<sup>2</sup>) |  |  |  |
| Protein | 57.39 | 109.49 | 63.31 |
| Ligand | 57.22 | 70.70 | 66.18 |
| R.m.s. deviations |  |  |  |
| Bond lengths (Å) | 0.004 | 0.004 | 0.004 |
| Bond angles (°) | 0.671 | 0.741 | 0.719 |
| Validation |  |  |  |
| MolProbity score | 1.65 | 1.8 | 1.73 |
| Clashscore | 7.01 | 7.82 | 7.24 |
| Poor rotamers (%) | 0.08 | 0.16 | 0.00 |
| Ramachandran plot |  |  |  |
| Favored (%) | 96.17 | 94.49 | 95.28 |
| Allowed (%) | 3.49 | 5.08 | 4.37 |
| Disallowed (%) | 0.34 | 0.44 | 0.34 |

